# Perturbation Variability Does Not Influence Implicit Sensorimotor Adaptation

**DOI:** 10.1101/2023.01.27.525949

**Authors:** Tianhe Wang, Guy Avraham, Jonathan S. Tsay, Sabrina J. Abram, Richard B. Ivry

## Abstract

Cerebellar-dependent implicit adaptation has been regarded as a rigid process that automatically operates in response to movement errors in order to keep the sensorimotor system calibrated. This hypothesis has been challenged by recent evidence suggesting flexibility in this learning process. One compelling line of evidence comes from work suggesting that this form of learning is context-dependent, with the rate of learning modulated by error history. Specifically, learning was attenuated in the presence of perturbations exhibiting high variance compared to when the perturbation is fixed. However, these findings are confounded by the fact that the adaptation system corrects for errors of different magnitudes in a non-linear manner, with the adaptive response increasing in a proportional manner to small errors and saturating to large errors. Through simulations, we show that this non-linear motor correction function is sufficient to explain the effect of perturbation variance without referring to an experience-dependent change in error sensitivity. Moreover, by controlling the distribution of errors experienced during training, we provide empirical evidence showing that there is no measurable effect of perturbation variance on implicit adaptation. As such, we argue that the evidence to date remains consistent with the rigidity assumption.

## Introduction

To achieve skilled movement, an ideal learner should track the state of the environment and adjust behavior accordingly. A skilled tennis player will adjust their anticipatory behavior more rapidly when facing an opponent who frequently switches from hitting crosscourt groundstrokes to lobs. Consistent with this hypothesis, experience modulates sensorimotor learning. For example, various studies have shown that the learning rate of the motor system is modulated by the rate of contextual change (Burge et al., 2008; Gonzalez Castro et al., 2014; Herzfeld et al., 2014).

However, the locus of context effects on motor learning remains unclear. Sensorimotor adaptation is driven by multiple learning processes (J. A. Taylor et al., 2014; Jordan A. Taylor & Ivry, 2014). Explicit mechanisms are critical for the volitional changes in behavior such as that required to adjust to a tennis opponent’s strategy. In contrast, the processes that keep the sensorimotor system precisely calibrated operate in an implicit manner. Accumulated evidence has suggested that while the explicit system can flexibly adjust to the environmental context, the implicit system is rigid, operating in a fixed manner that is insensitive to task demands (Jordan A. Taylor & Ivry, 03/2012). For example, the operation of the implicit system cannot be suppressed even when this results in an increase in performance error (Mazzoni & Krakauer, 2006; Morehead et al., 06/2017; Jordan A. Taylor & Ivry, 2014; Wang & Taylor, 2021; Wilterson & Taylor, 2021).

The characterization of the implicit adaptation system as rigid and context-independent has been challenged by studies in which the distribution of experienced errors is manipulated (S. T. Albert et al., 2021; Herzfeld et al., 2014). Albert and colleagues (S. T. Albert et al., 2021) conducted a series of visuomotor rotation experiments in which participants had to adapt to a 30° perturbation of the visual feedback. They compared conditions in which the size of the perturbation varied across trials (SD = 12°) or was constant (SD = 0°). Across the experiments, implicit adaptation was consistently attenuated in the high-variance condition. To account for this finding, the authors propose that the learning rate is modulated in a memory-dependent manner(S. T. Albert et al., 2021; Herzfeld et al., 2014). Specifically, sensitivity increases with repeated exposure to errors with the same sign and decreases when the sign reverses (Avraham et al., 2020; Hutter & Taylor, 2018). Such context-dependent effects have been observed with other learning processes where sensitivity is adjusted as a function of uncertainty (Behrens et al., 2007; Collins & Koechlin, 2012; Daniel M. Wolpert et al., 2003).

Although the Albert et al. study provides compelling support that performance is influenced by the variance of the perturbation, there is an alternative explanation for these results that does not entail memory-dependent modulation of the implicit system. The cornerstone for this alternative explanation is the observation that the implicit adaptation system exhibits a non-linear motor correction function: The correction response increases over a range of small errors up to the saturation point (around 7.5°), and eventually decreases for very large errors (Fig. 1a) (Bond & Taylor, 06/2015; Hutter & Taylor, 2018; Kim et al., 12/2018). A consequence of this non-linear function is that differences in the distribution of experienced errors as a function of perturbation variance may introduce a bias in terms of the elicited motor corrections. Thus, the differential response to high and low variance perturbations may not reflect flexibility within the adaptation system but be an emergent property of sampling across the non-linear motor correction function.

**Figure 1.**
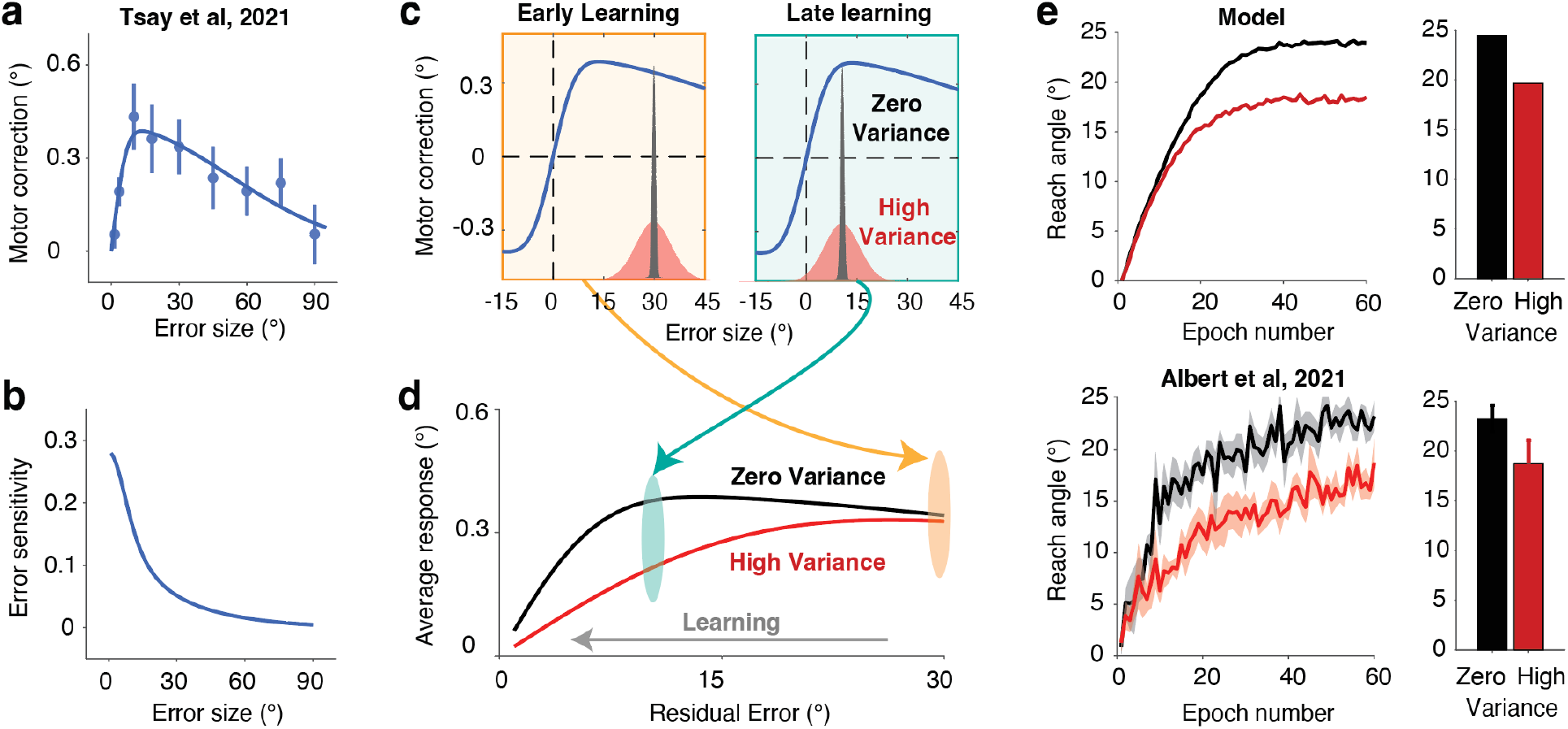
A non-linear motor correction function is sufficient to explain the effect of perturbation variability. **a**) Changes in reach angle from trial n to n + 1 as a function of the error size on trial n (Tsay et al., 2021). Dots represent group median values for each error size and bars represent the standard error of the means. We used these data to estimate the motor correction function given its relatively high sampling density. Solid line denotes the best-fitting model. **b**) Error sensitivity, defined as the motor correction divided by error size, reduces monotonically as error size increases. **c**) Perturbation variability impacts the distribution of experienced errors (red, high variability; black, low variability). This in turn will dictate the distribution of motor corrections. **d)** When the error is large at the start of learning (e.g., ∼30°), the average response is similar between the low- and high-variance conditions. However, as learning unfolds, the error distribution for the high-variance condition will span the concave region of the motor correction function (e.g., ∼10°), resulting in attenuation of the average motor correction. **e**) The learning function of implicit adaptation predicted by the non-linear motor correction (NLMC) model (top) provides a good approximation of the empirical results from Albert et al, Experiment 6 (S. T. Albert et al., 2021) (bottom). The right panels for each row show the reach angle late in learning. Error bars represent standard error. Shaded areas (left) and error bars (right) indicate standard error.

In the current study, we re-examine how perturbation variability influences implicit adaptation. We first take a modeling approach, using simulations to ask how variation in perturbation consistency will impact adaptation when sampling from a non-linear motor correction function. A core finding from the simulations is that as the variance of the perturbation increases, adaptation is attenuated due to sampling biases. Given that the simulations provide an alternative account of the results reported by Albert et al. (S. T. Albert et al., 2021), we then examine whether there is evidence of a memory-dependent change in error sensitivity in previous studies that randomized the perturbation sign and size across trials. The results of our re-analysis of these data sets also argue against the idea that the learning rate is modulated by error history. To complement these theoretical analyses, we conducted two new experiments in which we manipulated error variability while controlling for sampling biases between perturbations with high and low variance. The results show that the magnitude of adaption is independent of perturbation variability, further arguing against the hypothesis that implicit adaptation exhibits memory-dependent changes in error sensitivity.

## Results

### A non-linear motor correction function is sufficient to explain the effect of perturbation variability

Albert et al. provide evidence purporting to show that implicit adaptation is modulated by error history, a form of memory-dependent flexibility (S. T. Albert et al., 2021). Across a series of experiments, adaptation to a perturbation that varied in magnitude across trials (mean = 30° visuomotor rotation, SD = 12°) is weaker than adaptation to a fixed 30° rotation. To account for this finding, they proposed that the learning rate fluctuated with context, increasing in response to repeated errors and decreasing in response to errors that deviated from recently experienced errors. While they employed a variety of tasks, we focus here on their experiment (Exp 6) that isolates performance changes from implicit adaptation with minimal contribution from learning related to changes in explicit action selection (i.e., re-aiming).

Here we consider an alternative model that does not entail any experience-dependent modulation of the learning rate. The cornerstone of this model is the extensive literature showing that error sensitivity varies as a function of error size (Hutter & Taylor, 2018; Wei & Körding, 02/2009). When the function is described in terms of error rate (i.e., the change in performance as a proportion of the error magnitude), there is a monotonic decrease with error size (Fig. 1b). Alternatively, the function can be described in terms of the absolute size of the motor correction: When viewed this way, the function is non-monotonic, showing an increase in the size of the correction over a range of small errors, and then a reversal for larger errors (Fig. 1a). From a Bayesian inference perspective, this function captures the idea that large errors are discounted (Wei & Körding, 02/2009). Alternatively, this function may reflect the upper limits of plasticity in the sensorimotor system (Kim et al., 12/2018). Importantly, whether the function reflects a discounting process or limits in plasticity, the motor correction function is assumed to be a “primitive” or an established prior(Wei & Körding, 02/2009), existing independent of the recent context. We conducted a series of simulations to ask if a fixed non-linear motor correction function (NLMC model) can account for the core findings presented by Albert et al.

For our simulations, we used a motor correction function derived from a task in which the perturbation is randomized across trials (mean perturbation was zero) with relatively dense sampling across a large range of sizes (Fig. 1a)(Tsay, Avraham, et al., 2021). We examined the consequences of perturbation variability under the assumption that the motor correction function is fixed (Fig. 1c-d). Importantly, perturbation variability will impact the probability distribution of the experienced errors, and due to the nonlinearity of the motor correction function, will impact the average motor correction. If the experienced errors are all sampled from the descending linear zone, the average motor correction will be similar for perturbation conditions involving either low or high variability (Fig. 1c, yellow box). For adaptation to a visuomotor rotation of 30°, the errors during the early stages of learning will come from this part of the function. However, if the experienced errors span the concave region of this function—a situation that occurs when there has been some compensation for the rotation, the condition with high variability will result in an attenuated average response (Fig. 1c, green box). This attenuation occurs since the corrections to errors smaller than the mean error are no longer balanced by corrections to errors larger than the mean error.

Following the method of Experiment 6 in Albert et al., we simulated learning functions generated by the NLMC model for a visuomotor adaptation task using perturbations centered at 30° with either high variability (12° SD) or zero variability (0° SD). We used a classic state-space model with the trial-to-trial update determined by the motor correction function displayed in Fig 1a (Tsay, Avraham, et al., 2021) and the retention factor reported in Albert et al. (see Methods). The simulated results of the NLMC model capture the key features of their behavioral results: The high-variance condition results in a slower learning rate and larger residual error (i.e., lower asymptote, see Fig. 1e). Importantly, these simulation results do not depend on the specific shape of the motor correction function. The same pattern of results can be obtained using a wide range of functions in which the motor correction function either decreases or saturates for large errors (Fig. S2).

We note that one analysis reported by Albert et al. appears to be inconsistent with the NLMC model. The error sensitivity function is always higher for the zero-variance condition across different error sizes. However, additional simulations show that an estimate of the error sensitivity function cannot be recovered in experiments using a blocked design (e.g., mean = 30°, Fig. 2) due to the presence of motor noise (Fig. S3). Indeed, the recovered error sensitivity function is higher in the zero-variance condition than in the high-variance condition, even if we use the same error sensitivity function for both conditions. Thus, the higher error sensitivity estimated for the low-variance condition may be an analytic artifact.

**Figure 2.**
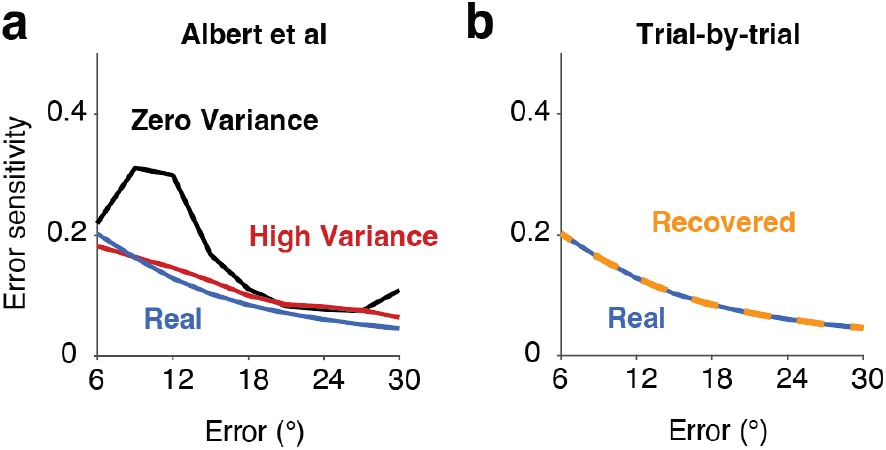
The error sensitivity function cannot be recovered in a block design. **a)** Recovered error sensitivity functions from simulations using a block design where the perturbation sign is constant. The Real function, based on the curve in Fig 1b, indicates the error sensitivity function that was used in the simulations. This function was fixed for high- and zero-variance conditions to generate the data. The estimated sensitivity functions for the two conditions fail to recover the original function. **b)** Recovered error sensitivity functions from simulations that used a trial-by-trial design where the direction and the size of the error were randomized across trials (mean perturbation = 0°). With this method, the recovered function matches the original function. Note that, by definition, a trial-by-trial design cannot have zero variance.

### Recent error history does not influence error sensitivity

Albert et al. (S. T. Albert et al., 2021) account for their behavioral results by reference to the Memory of Error model (MoE)(Herzfeld et al., 2014). This model assumes that the error sensitivity function is initially flat. During training, error sensitivity increases with repeated exposure to errors having the same sign and decreases when the sign of the error changes. The best way to conceptualize the consequences of this model is to consider the response to an error of the same size on trial n-1 and trial n+1 as a function of the error experienced on trial n (Fig 3a, left). Error sensitivity on trial n+1 will be higher than n-1 if the errors on trial n-1 and trial n have the same sign as; in contrast, error sensitivity on trial n+1 will be lower than n-1 if the errors on trial n-1 and trial n have opposite signs. Note, in blocked designs (e.g., Albert et al), participants mostly experience successive errors with the same sign; changes in error sign are relatively rare. As such, by the MoE model faster learning in the low variance condition is mainly because of sensitization after the same-sign errors. In the low variance condition, participants experienced errors of similar sizes successively so the enhancement of the error sensitivity accumulates across trials. In the high variance condition, the enhancement of error sensitivity is distributed across different error sizes and, as such, small for a given error size.

**Figure 3.**
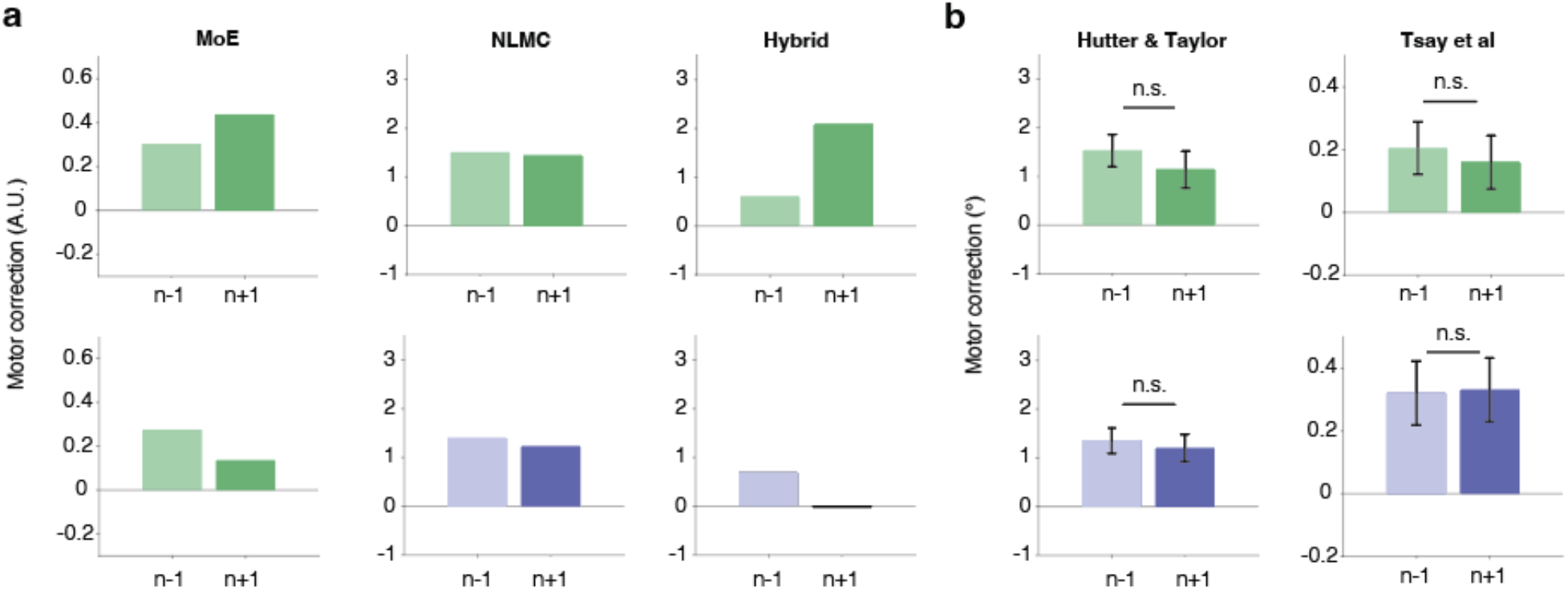
Data in trial-by-trial design does not support the Memory of Errors model. **a)** The effect of immediate error history on motor corrections as predicted by the MoE model (left), NLMC model (middle), and a hybrid model (right). For each model, we simulated data sets and then restricted the analysis to trial triplets in which the size of the errors on trials n-1 and n+1 had the same sign and were close in magnitude (difference <2°, see Methods). The results show the motor correction on these trials as a function of whether the sign of the error on trial n was the same (top) or different (bottom). The MoE and hybrid models predict a difference in motor correction, with an enhancement on trial n+1 when the sign is the same and an attenuation when the sign is different. The NLMC model predicts that the response will remain invariant. **b)** Behavioral results of a triplet analysis using data from two studies that used different methods to estimate changes in heading angle from implicit adaptation (left: Hutter and Taylor, 2018; right: Tsay et al, 2021). There was no evidence of an effect of recent error history in either data set. Error bars indicate S.E.

Given the biasing effects of motor noise, we turned to a trial-by-trial design in which the size of the perturbation was randomly selected on each trial (Mean = 0°, range: -90°∼90°). Frequent sign switches in the trial-by-trial design can also provide a strong test of whether there is any modulation of the error sensitivity. Figure 3a shows the results from these simulations for the MoE model (left) and NLMC model (middle). Consistent with the assumptions of each model, error history impacts the response on trial n+1 for the MoE model and has no effect on the NLMC model. To evaluate the models empirically, we re-analyzed the data from two published studies that used a trial-by-trial design to measure implicit adaptation (Hutter & Taylor, 2018; Tsay, Avraham, et al., 2021), focusing on trial triplets in which the size of the errors on trials n-1 and n+1 had the same sign and were close in magnitude (difference <2°, see method). Despite significant methodological differences between the two experiments, the pattern of results was similar: The relationship of the error sign between trial n-1 and trial n (same or different) had no effect on the motor correction on trial n+1 (Fig 3b). Thus, the re-analysis of these published data fails to support the MoE model, with no measurable effect of recent history on implicit adaptation.

In response to a preprint of this paper in which we described how the NLCM model can account for the effect of perturbation variability, Albert and Shadmehr (S. Albert & Shadmehr, 2022) proposed a hybrid model. Here they postulate that participants come to the experiment with a prior reflecting the non-linear motor correction function (Fig 1a), but that this function is modulated by the memory of errors. Nevertheless, this model also predicts a modulatory effect from recently experienced errors (Fig 3a, right) which is not supported by the trial-by-trial design. We elaborate on this model in the Supplementary Materials.

### Perturbation variability does not influence implicit adaptation

The previous simulations highlight two models that can account for the attenuation of adaptation when the perturbations are variable in a block design: 1) The MoE model which characterizes adaptation as a flexible process, sensitive to context, and 2) the NLCM model in which adaptation is rigid with the difference an emergent property of a sampling bias. The simulations highlight one differential prediction of the two models with the results failing to support the context effect predicted by the MoE model.

One limitation of the re-analyses is that we used data sets from experiments using a trial-by-trial design in which the perturbation had a mean of 0° and varied randomly from trial to trial. With this design, we cannot examine the cumulative effects of adaptation, a situation in which a change in learning rate would be especially pronounced. In this section, we present two experiments involving a block design, similar to that employed by Albert et al (S. T. Albert et al., 2021) but entail manipulations that control for the sampling bias problem.

For these experiments, we used an error clamp for the perturbation (Morehead et al., 06/2017). In this paradigm, the position of the feedback cursor is fixed with respect to the target location rather than the participant’s actual movement direction. Participants are instructed to ignore the feedback and aim directly to the target. Despite these instructions, their behavior shows all the hallmarks of adaptation: The hand path gradually shifts in the direction opposite the cursor and a pronounced aftereffect is observed when the perturbation is removed. Because the “error” is not reduced as adaptation occurs, the extent of adaptation is not yoked to the size of the error. Indeed, adaptation will asymptote around 20° -30° even in response to a clamped perturbation of 1.5° (Kim et al., 12/2018). Moreover, as mentioned before, the learning rate of this process scales with small errors before saturating in response to large errors (>7.5°), resulting in a non-linear function. One advantage of the clamped feedback over standard methods using response-contingent feedback is that there is no contamination from explicit learning processes (e.g., re-aiming); indeed, participants report their hand position to be close to the target throughout adaptation (Tsay et al., 2020).

In the first experiment, we used the error clamp method as an alternative way to test if adaptation is attenuated in a high variance condition compared to a no variance condition. For both conditions, the mean of the distribution was 12° and the size of the clamp was either drawn from a high variance distribution (SD=12°) or a distribution with zero variance (Fig 4a, top). We selected the mean value to be near the concave, non-linear zone of the response correction function. This will create error distributions similar to that experienced during the later phase of learning in Albert et al. where the difference between the high and zero conditions is greatest. By centering the distribution in the concave zone, we also maximize the sampling bias between the two conditions such that smaller and larger deviations from the mean value will not offset each other (keeping this constant across the experimental block). From this, the NLMC predicts a difference between the high variance and low variance condition (Fig. 4b, top). Note that theoretically, the MoE and the Hybrid model also predict a difference across two conditions. However, given the error sensitivity continuously increases while the error size never decreases, these models predict a very large asymptote (> 100°) using parameters estimated from the Albert et al. data.

**Figure 4.**
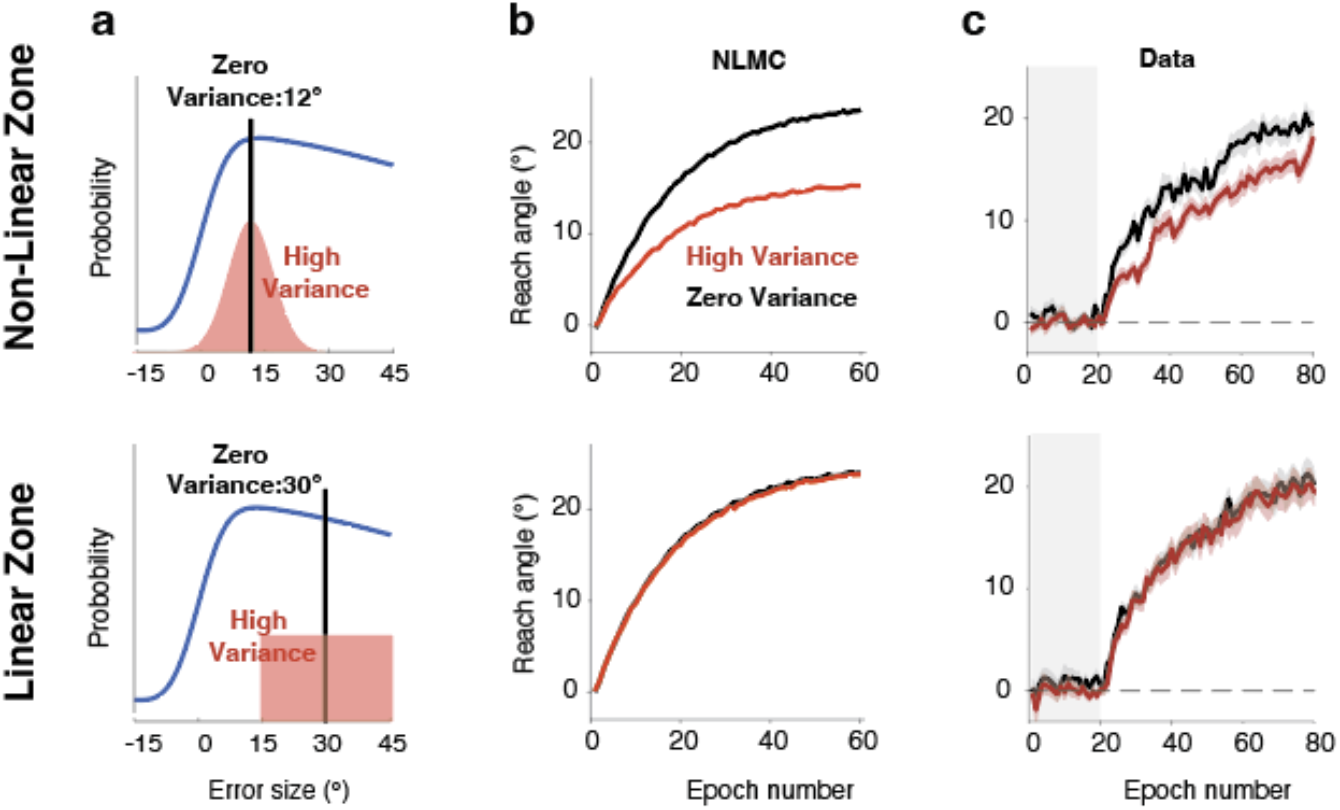
Influence of error variance on adaptation in the clamp rotation task. **a)** Distribution of clamp sizes in Exp 1 (top) and Exp 2 (bottom). In Exp 1, the mean was at 12° for both the High (red) and Zero (black) conditions, with the distribution for the High condition spanning the non-linear zone of the response correction function. In Exp 2, the mean was at 30° and the distribution for the High condition was restricted to the linear, descending zone of the response correction function. **b)** Simulations of adaptation functions based on the NLMC model. Note we do not visualize the prediction of the MoE and Hybrid models given that they predict asymptotic values greater than 100°. **c)** Experimental results in both experiments are consistent with the predictions of the NLMC model. Gray area indicates the no-feedback baseline. Shaded area indicates SE.

Similar to the prediction of NLMC, participants in the high variance group exhibited a slower rate and lower asymptote than participants in the zero-variance group (Fig. 4c, top). A mixed ANOVA showed a main effect of perturbation variance (F(1,67)=8.4, p=0.005, full results in table S1). Thus, a high variance perturbation results in attenuated implicit adaptation when the errors are sampled from the non-linear zone of the motor correction function.

The more critical test arises when the perturbation distribution is such that the sampled errors are restricted to one of the linear regions of the response correction function (Fig 4a, bottom). In Experiment 2, we opted to focus on large errors where the correction function is decreasing given that the ascending part is too narrow to adequately create a high variance condition. Rather than use a Gaussian distribution, we employed a uniform distribution for the High variance condition, with the clamp size ranging from 15° to 45°. In this way, we avoided trials in which the clamp was from the non-linear zone. For the Zero variance condition, the clamp was always at 30°. Both the MoE and Hybrid models predict that there will be an effect of perturbation variance, reflecting the slower sensitization of adaptation when there is a large difference in clamp size across neighboring trials. In contrast, the NLMC model predicts there will be no difference between the conditions. The results are again consistent with the NLMC model (Fig 4c, bottom). A mixed ANOVA showed no effect on perturbation variance (F(72,1)=0.01, p=0.92, full results in table S2).

We focused on the Hybrid model as an additional test of whether error sensitivity changes over the course of learning. Here we fit the data from both Exps 1 and 2 simultaneously, with the parameters determined at the group level. To estimate the distribution of the hyper learning rate and hyper retention rate of error sensitivity parameters, we fixed the other parameters and performed a bootstrapping analysis (Fig S5). The results showed that the hyper modulation rate was always zero (and the hyper retention rate of the error sensitivity was highly variable since it had no influence on the model’s behavior given the hyper learning rate is zero). Thus, this second test indicates that error sensitivity is not modulated during learning. Indeed, the best-fitting hybrid model becomes effectively, identical to the NLMC model.

Taken together, the results of the two experiments fail to support the hypothesis that adaptation is sensitive to error history. Rather, the data point to a model in which adaptation is a rigid process, one in which the output of the system is dictated by the size of the experienced error.

## Discussion

A hallmark of intelligence is the ability to modify behavior in response to an uncertain and variable environment. Consider the golfer out on the links on a blustery day. A skilled player will consider the direction and speed of the wind when choosing her club or orienting her stance prior to striking the ball. The unskilled player will also face this challenge, but her ability to learn from each outcome is likely to be limited given the uncertainty in determining if an errant shot was due to the impact of a wind gust or inconsistency in her swing.

Experimentally, numerous studies have highlighted how the rate of learning is modified as a function of uncertainty (Burge et al., 2008; Hsu et al., 2005; Körding & Wolpert, 2004; McGuire et al., 2014; Piray & Daw, 2021; Steyvers & Malmberg, 2003; Wei, 2010). For example, learning is attenuated when the size and direction of the perturbation randomly changes across trials compared to when the perturbation is fixed (Gonzalez Castro et al., 2014; Herzfeld et al., 2014). While it is clear that uncertainty influences how willing we are to change a policy or strategy (Behrens et al., 2007; Browning et al., 4/2015; Piray & Daw, 2021), the influence of uncertainty on the automatic, implicit process that keeps the sensorimotor system precisely calibrated remains the subject of debate. Many studies have shown that the implicit system is surprisingly rigid, insensitive to environmental statistics (Avraham et al., 2020; Hutter & Taylor, 2018) or task demands (Hutter & Taylor, 2018; Morehead et al., 06/2017; Jordan A. Taylor & Ivry, 2014).

In contrast, one influential theory, the Memory of Error model, posits that this system retains a history of experience memory, using this information to modulate error sensitivity (Herzfeld et al., 2014). Support for this model comes from a recent study reporting that the rate of adaptation is influenced by the variability of errors, and in particular is attenuated when exposed to an environment in which the perturbation schedules are variable compared to when it fixed (S. T. Albert et al., 2021). Here we report results that offer an alternative interpretation of the effect of error variability. In a series of simulations, we found that error variability impacts implicit adaptation because of a sampling bias present in a high variance condition compared to a low variance condition. This bias arises when one assumes that the trial-by-trial adjustment in the sensorimotor map is determined by a, fixed non-linear motor correction function: In the high variance condition, samples are drawn from both the ascending and descending (or linear) zones of this function, whereas in the zero variance condition, all of the samples are identical (subject to noise). The net result of this bias is that cumulative learning will be attenuated in the former. To complement the simulations, we conducted two experiments in which we controlled the sampling bias by using an error clamp. When the bias was present, we observed attenuated learning in a high variance condition compared to a low variance condition. Critically, when the bias was eliminated by limiting sampling to a linear zone of the motor correction function, there was no effect of error variability. Taken together, the simulations and empirical results are consistent with the hypothesis that implicit adaptation is rigid, insensitive to the uncertainty of the environment.

As noted above, our interpretation of the present work is based on the assumption that there exists, *a priori*, a fixed motor correction function. One could argue that the Memory of Error model provides a mechanistic account of how this non-linear function emerges with experience. Namely, error sensitivity will be lower for large errors simply because these are experienced less frequently than small errors. Indeed, as shown in Albert et al. (S. T. Albert et al., 2021), their model predicts a motor correction function that bears a non-linear shape within a limited range (0-30°; Fig. S4a), independent of how the function is initialized. We agree that this function is likely the result of experience. But this experience reflects a lifetime of experience rather than emergent from the specific conditions of a given experiment. This hypothesis is consistent with multiple lines of evidence. First, error sensitivity varies with error size in the initial responses to a perturbation(Bond & Taylor, 06/2015; Kim et al., 12/2018). Second, the non-linear motor correction function (and correspondingly, non-constant error sensitivity function) is also observed in tasks that randomly vary the sign and size of the perturbation from trial to trial (Hutter & Taylor, 2018; Robinson et al., 2003; Tsay, Avraham, et al., 2021; Wei & Körding, 02/2009). In contrast, the model presented in Albert et al. predicts a linear motor correction function (i.e., constant error sensitivity, see Fig. S4c) under a random perturbation schedule: The enhancing effect that would occur when an error of a given size is repeated would be nullified by the attenuating effect that occurs when the sign reverses for that sized error. Third, the motor correction function is not affected by the probability of sign reversals(Avraham et al., 2020; Hutter & Taylor, 2018). In sum, the assumption of an *a priori* non-linear motor correction function provides a parsimonious account of this prior work and the present results.

Given the astonishing flexibility with which humans learn and perform motor skills, it may seem paradoxical to argue that the implicit system is insensitive to environmental uncertainty. While the main basis of our argument is empirical— to date, the evidence points to an inflexible, rigid system--we note that this rigidity constraint may be a feature of a system that imposes some degree of modularity between processes associated with action selection and processes associated with movement implementation. The adaptation system is designed to keep the sensorimotor system exquisitely calibrated. While we, as experimenters can introduce large perturbations, these are unlikely to occur in natural behavior; rather, the adaptation system is designed to adjust to the small movement errors we typically experience due to misperceptions of the environment (de C. Hamilton et al., 2007; Flanagan et al., 1995; Odegaard et al., 2015; van Polanen & Davare, 2015), changes in bodily state (Burge et al., 2008; Johnston et al., 1998; J. L. Taylor et al., 2000), or intrinsic motor biases (Vindras et al., 1998, 2005). We adapt to these changes in an automatic manner, using error information to recalibrate the system within the present context without needing to change the control policy (i.e., action selection). In contrast, when operating in a novel environment (e.g., taking up a new skill), we are likely to experience large errors that may require the derivation of a new control policy. The ability to derive, implement, and evaluate a new control policy is key to the flexibility of the motor system. These processes are modulated by environmental uncertainty (Behrens et al., 2007; Browning et al., 4/2015). In essence, we propose that, whereas uncertainty can motivate a need for a change in action selection, the adaptation system is designed to ensure that the selected action is properly executed, independent of the optimality of that action.

We note that performance changes resulting from the adaptation system may be modulated by context, even if the parameters of this system are insensitive to environmental statistics(Heald et al., 2021). For example, when participants are re-exposed to a perturbation after the system has been reset (i.e., a washout phase), implicit adaptation is attenuated (Avraham et al., 2021). One account of this attenuation-upon-relearning phenomenon is that the system creates context-specific memories (Haruno et al., 2003; D. M. Wolpert & Kawato, 1998) which may be engaged simultaneously, resulting in interference. In this example, the washout phase would constitute a new context, one in which an opposing perturbation is used to reset the system or feedback is withheld to allow learning to decay. During relearning, performance will reflect the contribution of both contexts, with their weights determined by a generalization function. Assuming the memory developed during washout has a non-zero weight, the net effect will be slower adaptation during the relearning phase. Importantly, this effect occurs even though the error sensitivity of the system is not modulated by experience.

## Materials and Methods

### Models

#### Estimating the motor correction function

A primary goal of this study was to ask if the effect of error variability on adaptation might be a statistical artifact due to biases when sampling from a non-linear motor correction function. This function has been observed in many studies (Wei & Körding, 02/2009). For the simulations reported in this paper, we derived a motor correction function from data reported in previous studies that measured the trial-by-trial response of the implicit adaptation system to a range of error sizes. The primary data was from Tsay et al. (2021) since the perturbation sizes spanned a large range with relatively high sampling resolution. Non-contingent visual feedback was used in that study, a method designed to isolate implicit visuomotor adaptation(Morehead et al., 06/2017). The participant reached to a visual target and a feedback cursor was presented at the radial distance of the target but with an angular displacement defined relative to the target rather than the participant’s hand position. This displacement varied randomly across trials (19 conditions: 0°, ± 1.5°, 3.5°, 10°, 18°, 30°, 45°, 60°, 75°, 90°). Participants were fully informed that they had no control over the location of the feedback and should ignore it, always attempting to reach directly to the target. With these instructions, trial-by-trial changes in reaching behavior are implicit^12^.

To obtain a continuous motor correction function, we fit the empirical data with a model that encompasses features of two models for implicit learning: Visual uncertainty(Tsay, Avraham, et al., 2021) and relevance inference(Wei & Körding, 02/2009). We used the visual uncertainty model to capture the increase in the size of the motor correction as the error size increases over a range of small errors (Tsay et al. 2021). The model assumes that the average motor correction for a given error size depends on the probability, *p*(*positive* | *e*), that the participant perceived/considered the error in the correct direction:

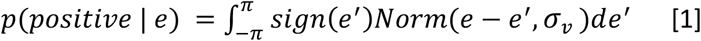

where *sign*(*e*^′^) indicates the sign of the perceived error, *e*^′^, and *σ*_*v*_ is the noise in the visual system; *Norm*(*e, σ*_*v*_)is the probability of observing *e* from a normal distribution centered at 0 with a standard deviation of *σ*_*v*_.We note that there are alternative accounts for the rise in the motor correction function for small errors(Hayashi et al., 2020; Kim et al., 12/2018; Makino et al., 2022). As such, our use here of the visual uncertainty model is motivated by pragmatic considerations; the results would be near-identical if we were to employ a different model (e.g., one based on uncertainty concerning motor noise).

The relevance estimation model was used to capture the decrease in the size of the motor correction for large errors. The core premise of this model is that large errors are likely to be attributed to sources independent of the motor system, and thus not relevant for recalibration of the motor system. Following the derivation of Wei and Kording (2009), error relevancy, *p*(*relevant* | *e*), is determined by the size of the error signal:

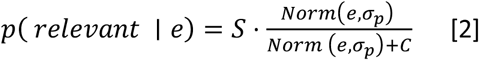

*S* and *C* are scaling factors that, jointly represent the prior belief of how likely an error is related to the motor system. *σ*_*p*_ is the proprioception noise.

Taken together, the motor correction function can be described as:

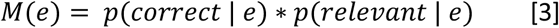

where *M*(*e*) is the motor correction to an error of size *e*. We obtained the continuous motor correction function by fitting this model with the median of the empirically observed error corrections(Tsay, Avraham, et al., 2021), minimizing the residual square error. We opted to use this relatively complex model because simple models (polynomial function, visual uncertainty model alone, relevance inference model alone) do not capture both the rapid rise and gradual decline of the motor correction function.

To calculate the error sensitivity function, *z*(*e*), the motor correction function was normalized by *e*:

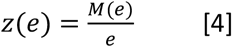

To construct the average motor correction function for the high-variance condition, *M*_*high*_(*e*), and low (zero) variance condition, *M*_*low*_(*e*), we computed the convolution between *M*(*e*^′^), and the perturbation distribution, *norm*(*e* − *e*^′^, *S. D*.), and integrated over *e*^′^:

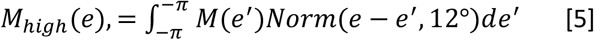

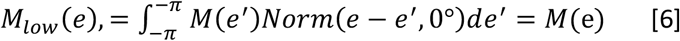

where *e* is the expectation value of *e*^′^. To simulate the results for Albert et al., we set the standard deviations to 12° and 0° in the High and Zero variance conditions, respectively.

To establish the generality of our findings, we repeated the simulations with a motor correction derived from the data in Hutter and Taylor (2018). In that study, the implicit component of learning was estimated by subtracting out the contribution of strategy use through verbal aim reports (Fig. S2 a-c). We performed additional simulations with a variety of hypothetical motor correction functions (Fig. S2 d).

#### State-space model

To model learning, we used a standard version of a state-space model:

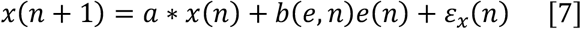

where *x* is the internal estimate of the motor state (i.e., hand movement required to compensate for the perturbation), *a* is the retention factor, *e*(*n*) is the size of the perturbation on trial *n, b* is the error sensitivity for a given error size, and *ε*_*x*_ represents planning noise.

The actual motor response on trial *n* is given as:

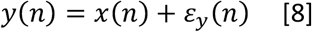

where *y* is the reaching direction relative to the target, determined by *x*(*n*) and the motor noise, *ε*_*y*_.

#### Non-Linear Motor-Correction Model (NLMC)

The NLMC model embeds the non-linear motor correction function into the state-space model. On each trial, error sensitivity is directly read out from the motor correction function based on the experienced error:

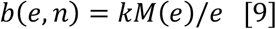

where *k* is a constant that scales the error sensitivity function (see details below). Note that whereas error sensitivity function, *b*, is modulated by experience in the MoE model; it is fixed across trials in the NLMC model. So that we can rewrite the Eq[7] as follow:

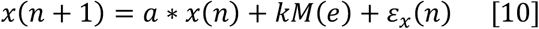

#### Memory of Error Model (MoE)

We compared the NLMC model to the MoE model introduced by Herzfeld (Herzfeld et al., 2014) and extended by Albert (S. T. Albert et al., 2021). In the MoE, error sensitivity, *b*, is a constant at the beginning of learning:

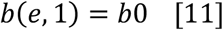

The key assumption of the MoE model is that *b* is modulated by the experienced error during training. Specifically, *b(e,n)* increases when *e(n-1)* and *e(n)* share the same sign, and *b(e, n)* decreases when the sign for *e(n-1)* and *e(n)* differ:

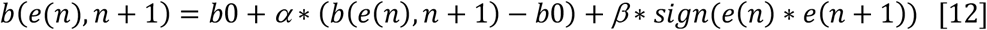

where *β* and *α* are the learning rate and retention rate of *b*, respectively, and *b*(*e*) is binned within every 3°. Note, we use the 3° bin to keep it consistent with (S. Albert & Shadmehr, 2022). The performance of the model won’t be qualifiedly different when changing the size of the bin or replacing the bin with a Gaussian Gain.

#### Hybrid model

Motivated by a commentary provided by Albert and Shadmehr(S. Albert & Shadmehr, 2022), we included a model that combines core features of the NLMC and MoE models. This model initializes error sensitivity according to the non-linear motor correction function and modulates the sensitivity values following the algorithm of the MoE model:

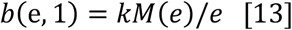

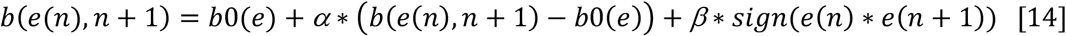

#### Model simulations

For simulations reported in the main text, we simulated the results of Albert et al.(S. T. Albert et al., 2021) with the NLMC model using a motor correction function, *M*(*e*), fit with the data from Tsay et al (2021, see Fig. 1). In a secondary analysis, we repeated this process using the data from Hutter and Taylor (2018, see Fig. S2) to obtain the motor correction function. While the shapes of those functions are similar, the overall magnitude of the motor correction differs between the studies. This is not surprising since the experimental apparatus and methods used to measure implicit adaptation had significant differences. For example, whereas Tsay et al. used endpoint feedback, Hutter and Taylor used continuous feedback (Hutter & Taylor, 2018; Tsay, Avraham, et al., 2021). Adaptation is attenuated with the endpoint feedback. For this reason, we scaled the motor correction function such that sensitivity to a 30° error (the initial error experienced when the perturbation is introduced in Albert et al.) was the same for the simulations using the two estimates of the motor correction function. This was done by setting *k* = 5 for the function derived from Tsay et al. and *k* = 0.4 for the function derived from Hutter and Taylor. The retention factor *a* was set as 0.945, a value obtained from the washout phase of Experiment 6 in Albert et al. For simulations with the MoE, we used the parameters reported by Albert et al.: *α* = 0.9568, *β* = 0.0558, *b*0 = 0.037, *a* = 0.945.

We assume that learning is the same for reaches to each of the four targets and thus only simulated the learning of one target for each participant. We set *ε*_*x*_ as 0 for simplicity since it does not have any influence on learning. The motor noise, *ε*_*y*_ were sampled from normal distributions with a mean of 0 and an S.D. of 3°. We note that for motor variability, we only include motor noise measured from the baseline section. This might underestimate the motor variability since the motor variance for the individual participant during the learning is much larger compared with the prediction of the model, perhaps because of active exploration. Consistent with the experimental design of Albert et al., the perturbation in the zero-variance condition was always set to 30°. The perturbations in the high-variance condition were randomly sampled from a normal distribution with a mean of 30° and S.D. of 12°. The displayed simulation results were the average of the simulated behavior of 1000 participants for each perturbation variance condition.

To simulate the model predictions in the trial-by-trial design (Hutter & Taylor, 2018), we fixed all the parameters of the model as we described above and applied a different perturbation schedule. The motor correction function was from Tsay et al., 2021. The errors were sampled from 0°, ± 4°, 8°, 16°, 32°. Each error was visited once before being repeated. The displayed simulation results were the average of the simulated behavior of 1000 participants that performed 200 trials. To simulate the model predictions of the NLMC model for our error-clamp rotation experiments, we applied the same parameter set as used in the previous simulation.

#### Model fitting

To fit the Hybrid model to Albert et al., we applied a grid search in which *α* and *β* ranged from 0 – 1 with a step size of 0.01, and *k* ranged from 0 - 5 with a step size of 0.1. We fit the model to the learning functions from the High and Zero conditions simultaneously. Given that there is minimal generalization between targets spaced by 90°(Morehead et al., 06/2017), we simulated learning at each target separately, repeating this process for each participant. We used the actual errors experienced by the participants as input to the model in generating the learning functions. To evaluate the fit, we compared the behavioral data with the model prediction to compute the squared residual error. Given that we employed the actually experienced errors, motor noise was set to 0 during model fitting. The squared residual errors were summed across targets and participants. In describing the performance of the Hybrid model, we use the parameter set that generated the smallest sum squared residual error. We also fitted the hybrid model to Exp 1-2 simultaneously. Besides the hyper parameters *α, β*, and *k*, we also fitted the four parameters that decided the non-linear motor correction function (*σ*_*v*_, *σ*_*p*_, *S, C*). After that, we fixed *σ*_*v*_, *σ*_*p*_, *S, C*, and *k*, and performed the bootstrap at the participant level for 100 times to estimate the variation of *α, β*.

### Albert et al., 2021

Albert et al. examined the influence of perturbation variability on sensorimotor adaptation. Across several experiments, they found learning with high perturbation variance is attenuated compared to the no variance condition, and when they isolated the contribution of the implicit and the explicit systems, they found that this difference rests exclusively in the implicit system. We focus here on Experiment 6 of their paper since this used a design in which performance changes are likely to be dominated by implicit adaptation. Moreover, this is the experiment they used to formally examine the MoE model. To limit the use of explicit strategies in that experiment, participants were required to initiate the response within 250 ms of the onset of the target, minimizing preparation time. The target appeared at one of four locations (+/-45°, +/-135°), with each location visited once during a cycle of four trials. On perturbation trials, the visual feedback was rotated relative to hand position. The rotation was always 30° in the Zero variance condition; in the High variance condition, the actual rotation on each trial was sampled from a normal distribution with mean = 30° and SD = 12°. The experiment consisted of 40 baseline trials, 240 training trials with perturbations, and 40 no-feedback trials.

### Analyses of data in trial-by-trial designs

As noted, MoE postulates that error sensitivity will increase when recently experienced errors are similar and decrease when they are different. However, flip of error rarely occurs in blocked designs so the primary effect of error memory is to enhance familiar errors. In contrast, trial-by-trial designs in which the size of the perturbation varies randomly across trials will yield many sign changes, providing a strong test of the MoE mode.

To determine whether the error sensitivity is modulated in a trial-by-trial design, we performed an analysis of published data from two studies (Hutter and Taylor, 2018 and Tsay et al., 2021). The MoE and Hybrid models postulate that error sensitivity will increase when the errors experienced on trials n-1 and n are similar: If a similar error is experienced on trial n+1, the change in heading angle will be larger than that which had been observed on trial n-1. Likewise, if there is a sign change between trials n-1 and n and the error on trial n+1 is similar to that of trial n-1, the response to the former will be attenuated with respect to the response to the latter. To examine this prediction, we selected triplets of trials where the errors on trials n-1 and n+1 were of the same sign and the difference between them was smaller than 2°. We used a criterion of 2° boundary to maximize resolution while maintaining a reasonable sample size (∼20 trials/participants). Following the implementation of the MoE model in Herzfeld et al. (XX), we computed the change in hand angle on trial n (relative to n-1) and n+2 (relative to n+1) respectively, grouping the results based on whether the error on trial n shared the same sign as n-1 and n+1. Note, when all of the trials in a triplet share the same error sign, learning will accumulate so that the change in hand angle will reflect single-trial learning as well as forgetting. Thus, we fit a state space model to estimate the retention rate of the system and corrected for forgetting in calculating the change in heading angle. We performed an identical analysis on simulated data generated from each of the three models.

### Clamp rotation task

We report two experiments using a blocked design with clamped feedback. A challenge to examining how perturbation of variance influences implicit adaptation is that the effect of perturbation variability will be impacted by the non-linear motor correction function. By making the error independent of the movement, we can control the distribution of experienced errors and thus control the potential effect of sampling bias due to the non-response motor correction function.

#### Participants

160 young adults (76 female, age: 24.3 + 6.5) were recruited for online experiments using the website Prolific.io. The participants performed the experiment on their personal computers through OnPoint, a web-based platform for motor learning experiments(Tsay, Ivry, et al., 2021). Based on a prescreening survey employed by Prolific, the participants were right-handed and had normal or corrected-to-normal visions. The participants provided informed consent as part of a protocol approved by the Institutional Review Board of the University of California Berkeley. Participants received $8 financial compensation for their participation.

#### Experimental task and protocol

To start each trial, the participant moved a white cursor (radius: 0.6% of screen height) to a white start circle (radius: 1% of the screen height) positioned at the center of the screen. After 500 ms, a blue target circle (radius: 1% of the screen height) appeared with the radial distance set to 40% of the screen size. There were four possible target locations (+/-45°, +/-135°). The participant was instructed to produce a rapid, out-and-back movement, attempting to intersect the target. In most trials (see below), the cursor was visible during the outbound phase of the movement until the amplitude of the cursor movement reached the target distance. The cursor was then presented at the target distance for 50ms, after which both the cursor and target disappeared. To help guide the participant back to the start location, the cursor appeared when the hand was within 40% of the target distance. If the movement time was >500 ms, the message “Too Slow” was presented on the screen for 500 ms.

The experiment began with a familiarization phase that consisted of two blocks of trials: 40 without feedback (cursor blanked at movement onset and reappeared only when the hand was within 40% of target distance on inbound movement) followed by 40 trials with veridical, online feedback where the cursor tracked the position of the hand throughout the outbound of the movement. Following this, participants were tested on a learning block consisting of 240 trials with clamped feedback. For these trials, the radial position of the cursor was based on the radial extent of the movement, but the angular position was predetermined with respect to the target position (see below) and followed a straight-line trajectory along this angle. Thus, the angular position was independent of the position of the hand. The cursor remained visible until the movement reach the radial distance of the target and then disappeared. Prior to the learning phase, participants were informed that they would no longer control the movement direction of the cursor. They were instructed to ignore the cursor, attempting to always reach directly to the target.

To examine the effect of perturbation variance on adaptation, we conducted two experiments, one in which the clamp angle was always sampled from the non-linear zone of the motor correction function (Exp 1) and the other in which the clamp angle was always sampled from a linear zone of the motor correction function (Exp 2). We compared the effect of Zero and High variance clamps. In Exp 1, the clamp angle was fixed at 12° for the Zero variance group whereas the clamp value on each trial in the High variance group was sampled from a Gaussian distribution centered at 12° and a standard deviation of 12°. As such, participants will experience errors from the ascending and descending zones of the motor correction function. In Exp 2, the clamp angle was fixed at 30° for the Zero variance group and the clamp angle for each trial in the High variance group was sampled from a uniform distribution that ranged from 15 to 45°. We opted to use this distribution to ensure that all of the experienced errors were from the descending zone of the motor correction function.

#### Data analyses

Data analyses were conducted with MATLAB 2020b. Hand angle was calculated as the angular difference between the target and the hand position at the target radius. We averaged the hand angle values over cycles of four trials (1 reach to each of the four targets per cycle). Positive values indicate hand angles in the opposite direction of the perturbation, the direction one would expect due to adaptation. We excluded trials with a movement time that was longer than 500 ms or an angular error larger than 70°. We excluded the data from six participants who had less than 70% valid trials, resulting in final sample sizes of 76 for Exp 1 (Zero=37; High=39) and 78 for Exp 2 (Zero=38; High=40)

To examine the effect of clamp variance in the linear-zone and non-linear zone, we used a mixed ANOVA. Epoch number, error variance, and their interaction were treated as fixed effects and participant as a random effect. The dependent variable was hand angle.

## Data availability

The data and code that used in this project are available at https://osf.io/exg4c/

## Author contributions

All of the authors contributed to the conceptual development of this project. T.W. performed the experiments, analyzed the data, and wrote the initial draft of the paper, with all of the authors involved in the editing process.

## Supplementary Information

## Supplementary Results

### The influence of motor variability on estimation of error sensitivity

Consistent with their Memory of Error model (MoE), Albert et al. (2021) report that error sensitivity is higher in the zero variance condition compared to the high variance condition (see Figure 4a in their paper). Superficially, this result appears to be at odd with the Non-Linear Motor-Correction model (NLMC). However, as reported in the main results, our simulations indicate that an estimate of the error sensitivity function cannot be recovered from the block design used in their study. More generally, we believe it is difficult to recover an error sensitivity function in designs in which the effects of learning accumulate. We develop the theoretical basis of this claim in this section.

Error sensitivity for a given trial n, with error size, *e* is calculated as follows:

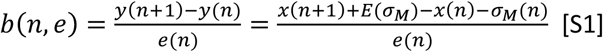

where *y*(*n*) is the movement angle, and *x*(*n*) is the state of the internal model on trial n. Consider the situation late in learning where performance is at asymptote with the state largely unchanged from trial to trial (*x*(*n* + 1) = *x*(*n*)). When the perturbation is fixed, variation in heading angle is dominated by motor noise (*σ*_*M*_). Given that the average motor noise by definition is 0, (*E*(*σ*_*M*_)=0), we have

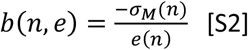

Thus, error sensitivity, *b*(*e*), when estimated this way would not reflect the real error sensitivity of the adaptation system. Rather it is function determined by error size and the motor noise for that trial. Given that the average residual error (*RE*) at asymptote is a constant, we can rewrite S2 as:

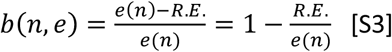

From S3 we see that *b*(*e*) does not reflect the error sensitivity of the system; rather, the estimate of *b*(*e*) increases as the error size decreases when estimated at asymptote.

We note that the above example comes from the case where learning is saturated. However, the estimate of error sensitivity will be contaminated by this problem at all phases of learning in a block design. It will be especially pronounced when the hand angle approaches the asymptote or when motor noise is large (Fig. S5). The problem is not present in a trial-by-trial design where the mean perturbation is 0° since there will be no accumulated learning. As such, error sensitivity can be reliably estimated (confirmed in our simulations, see Fig 2b).

### A hybrid model in the implicit system

In response to a preprint we posted that described how the NLCM model provides an alternative account of the effect of perturbation variability, Albert et al. proposed a hybrid model in which baseline error sensitivity follows the non-linear motor correction function (a prior) but is modulated by the memory of errors(S. Albert & Shadmehr, 2022). This model can predict the large difference between high and zero variance conditions observed early in learning (e.g., around epoch 10), an effect that is not captured by either the NLMC or MoE models (Fig. S1a).

However, there are some serious limitations with a hybrid model of this form. First, to explain the large difference during early learning, the best-fitted hybrid model predicts a very large hyper learning rate of the error sensitivity (*a*), which is 0.21 (Fig S1b). Given that the model fit results in estimated baseline error sensitivity of 0.04, the model assumes a five-fold increase in error sensitivity after one trial, a rate of changes that seems unreasonably fast (Fig S1c). Moreover, the hybrid model predicts a hyper retention rate (*b*) of 0.19. This would suggest that the change in error sensitivity will only last for a single trial and, thus, result in large instability in error sensitivity across trials (Fig S1c). Similarly, the Hybrid model predicts a marked change of error sensitivity across trials in a trial-by-trial design (Fig 3a). These predictions fail to conform with the empirical results showing no change in error sensitivity across trials in all conditions.

We note that the large effects on performance that are observed early in learning are sometimes due to the deployment of an explicit strategy (J. A. Taylor et al., 2014). To minimize strategy use, Albert et al. restricted preparation time. However, this manipulation may not completely block the involvement of a re-aiming strategy (Haith et al., 2015; Huberdeau et al., 2019), and this issue is likely more relevant in the Zero variance condition than in the High variance condition. Explicit contributions to learning might be more prominent in the former because the evaluation of an aiming strategy would be easier given the fixed perturbation size. Indirect support for this hypothesis comes from the data reported in Exps 1-3 of Albert et al. In these experiments, there was no limit on preparation time and performance changes can be assumed to reflect contributions from implicit and explicit processes. Under these conditions, the difference between the Zero and High variance conditions only emerges when performance is near asymptote. We note that this is predicted by the NLMC model since the sampled error at asymptote will span the non-linear zone.

**Fig. S1.**
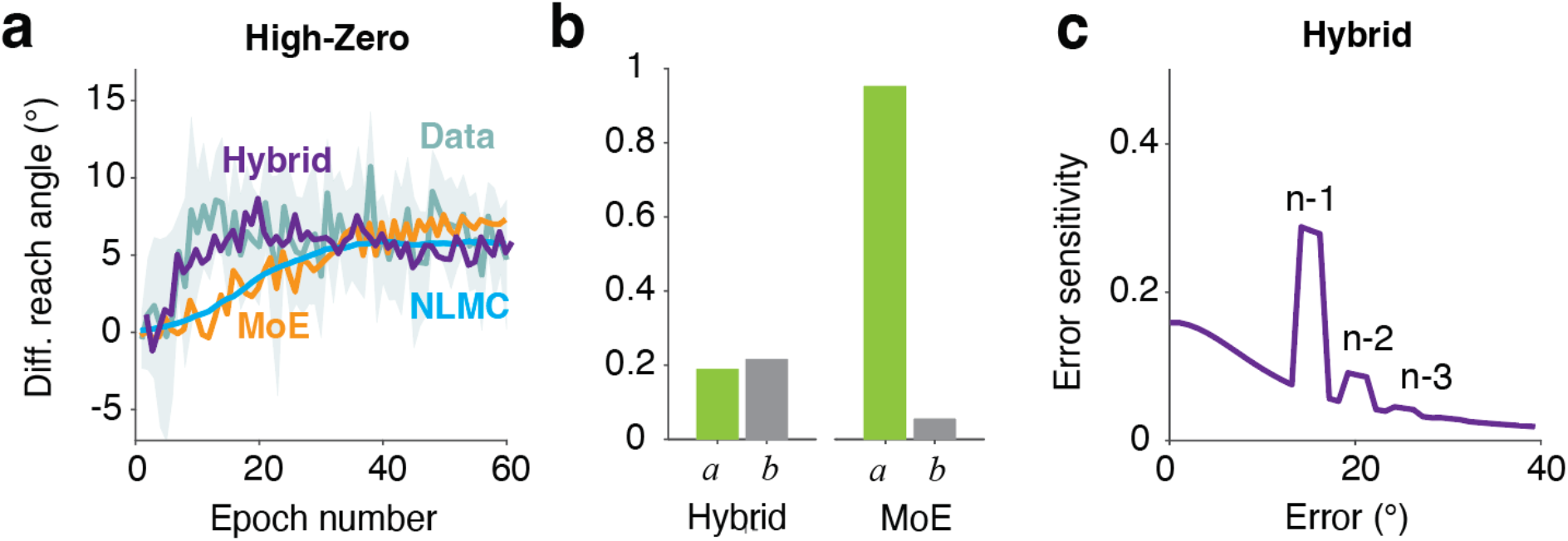
Evaluation of a Hybrid model for implicit adaptation. **a)** The difference between hand angle for the Zero and High variance conditions predicted by the MoE, NLMC, and Hybrid models, along with the data from Exp 6 of Albert et al (Shaded area indicates S.E.) In the actual data, the difference peaks early in the learning block (around epoch 10). Only the Hybrid model is able to capture this effect. **b)** However, this is achieved by having a large hyper learning rate (*b*) and small hyper retention rate (*a*, 1-forgetting) for error sensitivity. Not only are these parameter values unrealistic but they are quite different from those estimated for the MoE model. **c)** The hybrid model predicts a fast change of error sensitivity across trials. In this example, the experienced errors for the first five trials are 30°, 25°, 20°, 15°, and 10°. The error sensitivity calculated for n-1 is much higher than the baseline level because of the high learning rate for error sensitivity (*b*). This effect does not persist when sensitivity Is calculated for trials n-2 and n-3 since the retention rate of error sensitivity is very small.

**Fig. S2.**
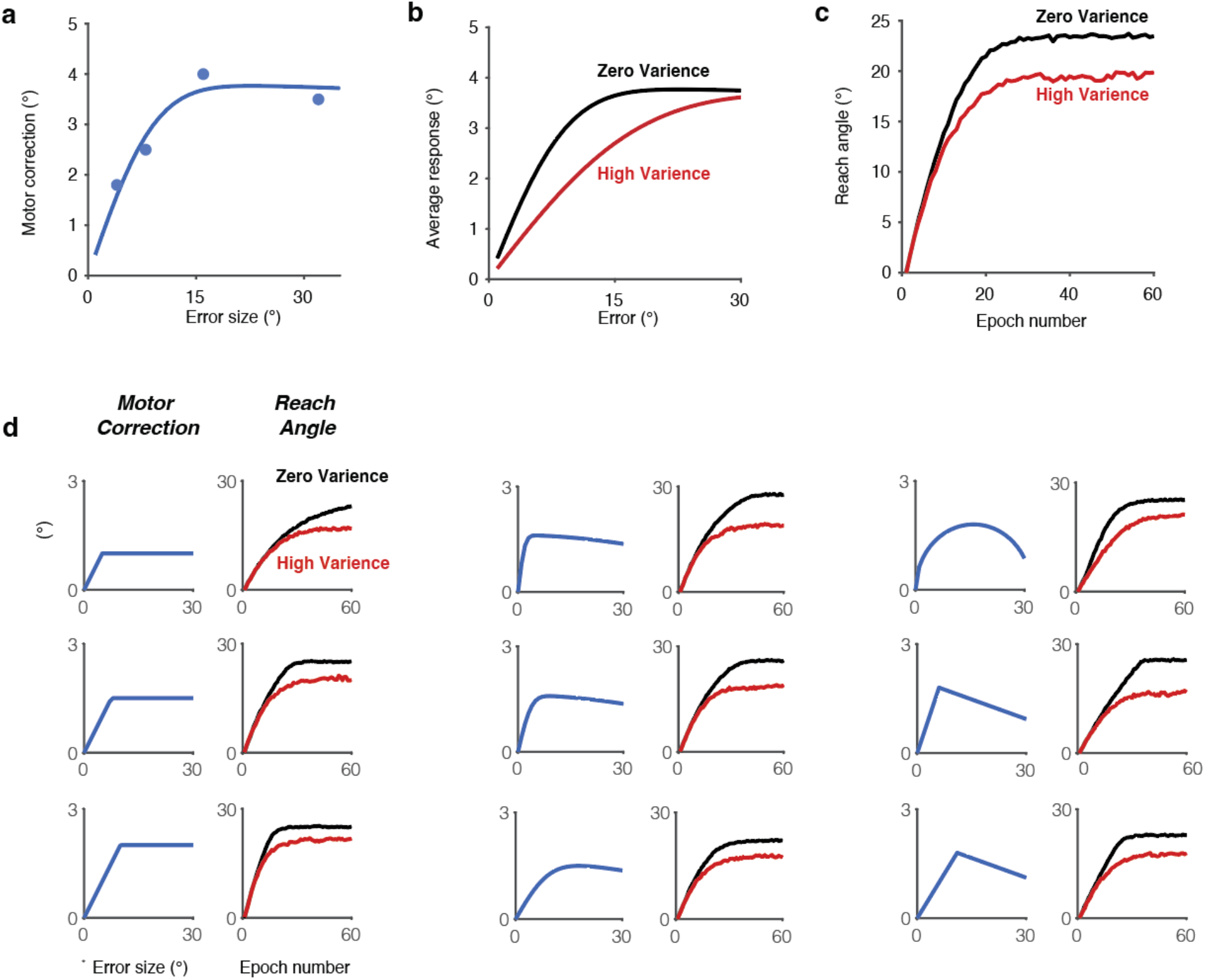
NLMC model captures the effect of error variability when using motor correction function derived from Hutter and Taylor (2018) or using hypothetical motor correction functions. **a)** Size of the motor correction as a function of the error experienced on the previous trial. Dots correspond to sampled values in Exp. 2 of Hutter & Taylor. Solid line indicates the best-fit model. **b**) Average motor correction as a function of mean error size under low (black) and high (red) perturbation variability conditions. **c)** Simulations of the NLMC model showing estimate of implicit adaptation. **d)** Effect of perturbation variability holds across a range of hypothetical motor correction functions. For each subplot, the motor correction function is depicted on the left and simulations of the learning functions for High and Zero variance conditions are shown on the right. In all of the examples, the hypothetical motor correction function increases in response to small errors and either saturates (left column) or decreases in response to large errors (middle and right columns). In each case, adaptation is attenuated in the high variability condition (red) relative to the zero variance condition (black).

**Fig. S3.**
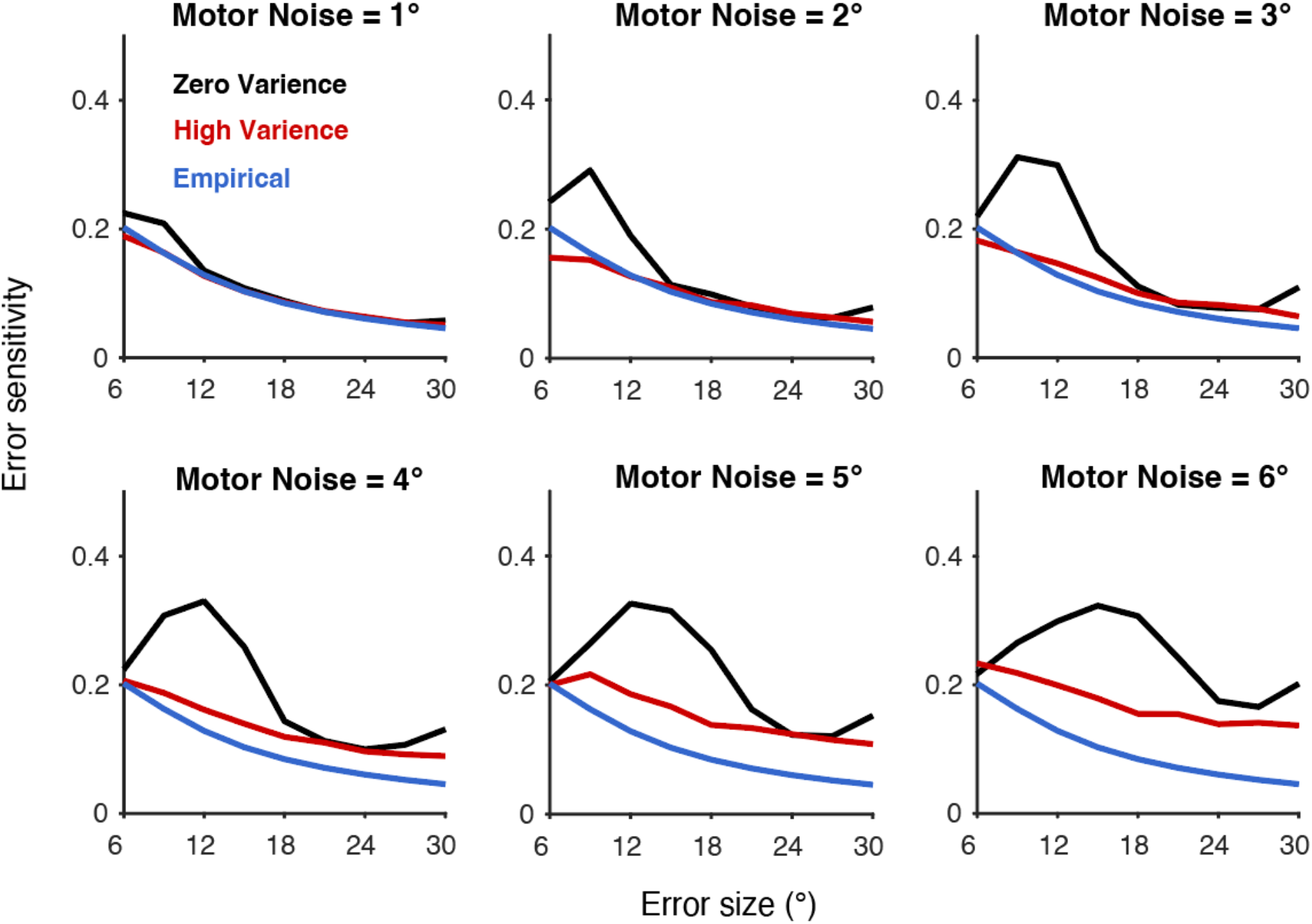
The error sensitivity function is not recoverable from data obtained in a blocked design experiment where adaptation will accumulate across trials. The six panels depict the effects for different levels of motor noise (range: 1-6°). Note that the recovered error sensitivity function is always higher in the Zero condition.

**Fig. S4.**
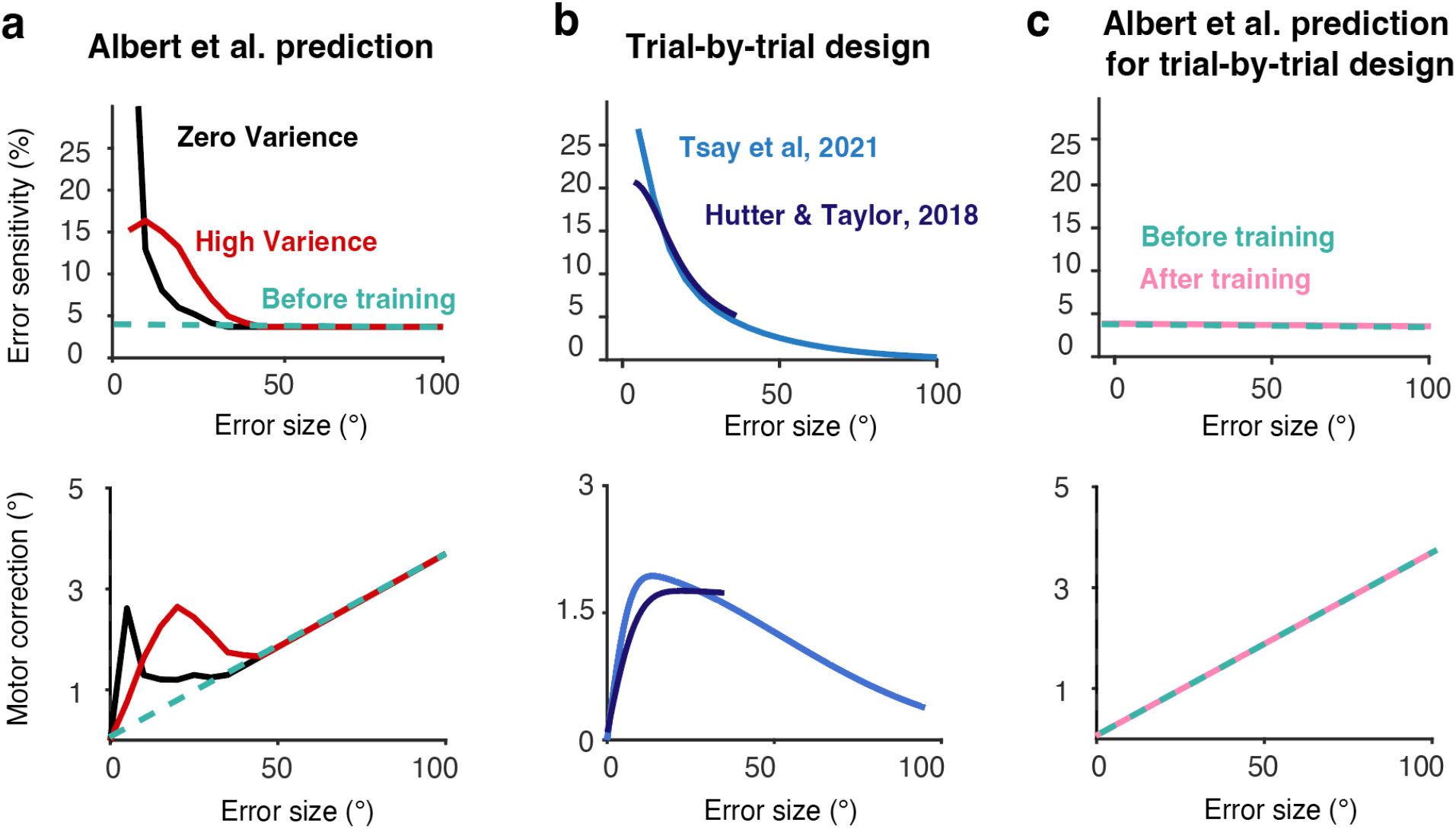
The Memory of Errors model fails to predict the error sensitivity function when the perturbation size varies randomly from trial to trial (mean = 0°). **a)** Albert et al. assumed that the error sensitivity function is initially a constant (top), resulting in a linear motor correction function (bottom). By itself, this function fails to capture the effect of perturbation variability. In their model, experiencing consistent or inconsistent errors will respectively, increase or decrease error sensitivity. This memory process will result in a non-constant error sensitivity function (top) and a non-linear motor correction function (bottom) for error sizes that were experienced during the experiment (0 to 30°). **b**) Error sensitivity function (top) and the non-linear motor correction function (bottom) estimated from the trial-by-trial designs used by Tsay et al (2021) and Hutter and Taylor (2018). **c**) In the trial-by-trial design, the MoE predicts that the shape of the error sensitivity function will be maintained over the course of learning given that errors of the same sign and opposite signs will result in no net change in sensitivity. The model fails to match the sensitivity function (top) or motor correction function (bottom).

**Fig. S5.**
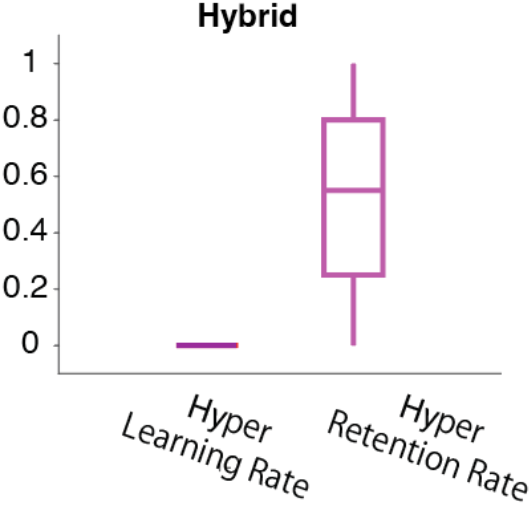
Model fitting shows that the learning rate is not modulated during the learning phase of Experiments 1 and 2. **a)** We fit the hybrid model to the bootstrapped (100 times) data set generated from Exps 1-2. The best fitted hyper learning rate for error sensitivity was always zero. As a consequence, the hyper retention rate of the error sensitivity was highly variable since it had no influence on the model behaviors given the hyper learning rate is zero.

**Table S1:** Results of the mixed ANOVA of EXP 1

|  | F | P |
| --- | --- | --- |
| Epoch | F(129,8643)=188.4 | <0.001 |
| Variance | F(1,67)=8.4 | 0.005 |
| Epoch* Variance | F(129,8643)=1.7 | 0.07 |

**Table S2:** Results of the mixed ANOVA of EXP 2

|  | F | P |
| --- | --- | --- |
| Epoch | F(129,9288)=156.1 | <0.001 |
| Variance | F(1,72)=0.01 | 0.92 |
| Epoch* Variance | F(129,9288)=0.7 | 1.0 |

